# A Sub-Minute Resolution Prediction of Brain Temperature Based on Sleep-Wake State in the Mouse

**DOI:** 10.1101/2020.08.12.246405

**Authors:** Yaniv Sela, Marieke M.B. Hoekstra, Paul Franken

## Abstract

While brain temperature is of neurobiological and clinical importance, it is still unclear which factors contribute to its daily dynamics and to what degree. We recorded cortical temperature in mice alongside sleep-wake state during 4 days including a 6h sleep deprivation, and developed a mathematical tool to simulate temperature based on the sleep-wake sequence. The model estimated temperature with remarkable precision accounting for 91% of its variance based on three main factors with the sleep-wake sequence accounting for most of the variance (74%) and time-of-day (‘circadian’) the least (9%). As third factor, prior wake prevalence, was discovered to up-regulate temperature, explaining 43% of the variance. With similar accuracy the model predicted cortical temperature in a second, independent cohort using the parameters optimized for the first. Our model corroborates the profound influence of sleep-wake state on brain temperature, and can help differentiate thermoregulatory from sleep-wake driven effects in experiments affecting both.

## Introduction

Brain temperature is a fundamental variable capable of affecting numerous neural processes, from basic properties such as nerve conduction velocity, passive membrane potential, and synaptic transmission, to global regulation of brain activity (Wang et al., 2014). At the same time, neuronal activity is one of the main determinants of brain temperature (Kiyatkin et al., 2002), showing that brain activity both affects and is affected by fluctuations in temperature. Clinically, brain temperature is known to increase during a large number of common pathological conditions such as stroke or head injury (Mrozek et al., 2012), and lowering temperature is used as clinical intervention to protect the brain during recovery from hypoxic events (Faridar et al., 2011). Since heat plays a crucial role in neuronal functioning (Kiyatkin, 2010; Alonso and Marder, 2020) and brain tissue is very sensitive to thermal damage (Yarmolenko et al., 2011), it is important to understand which main factors normally contribute to brain temperature.

The sleep-wake states, Non-Rapid-Eye-Movement (NREM) sleep, REM sleep, and wakefulness, define brain states associated with distinct neuronal activities and metabolism (Nir et al., 2013). Accordingly, the latter two (active) brain states are accompanied by increases in brain temperature, while NREM sleep by a decrease (Hayward and Baker, 1969; Obal et al., 1985; Franken et al., 1992a; Hoekstra et al., 2019). By quantifying the relationship between sleep-wake state and brain temperature we found that the sleep-wake distribution explains 84% of variance of brain temperature in the rat (Franken et al., 1992b), a finding we recently replicated in the mouse (Hoekstra et al., 2019). However, the early analysis has been criticized for overestimating the impact of sleep-wake state due to averaging over hourly intervals and ignoring other contributing factors such as locomotion (Heller et al., 2011), and to the sequential nature of assessing the contribution of different factors such as circadian influences (Witting and Mirmiran, 1997). Although recent studies found the contribution of locomotor activity to brain temperature to be small (Shirey et al., 2015; Hoekstra et al., 2019) and confirmed that circadian factors do not importantly contribute to brain temperature (Baker et al., 2005), the underlying concerns for using hourly values and fixed order of assessing the importance of contributing factors were not directly addressed.

To resolve these outstanding issues, we used a recent dataset (Hoekstra et al., 2019) to develop a mathematical model that simulates changes in brain temperature in a dynamic context and at the high time resolution required to account for its rapid fluctuations at sleep-wake state transitions. By going beyond our previous models we succeeded in explaining 91% of the variance in brain temperate and to reduce the model error to 0.26°C out of a dynamic range of 3.13°C. In addition to accurately capturing the short-term dynamics associated with sleep-wake transitions, the model revealed longer-term wake prevalence as a novel factor altering the range of values at which brain temperature is regulated. The third, circadian factor, explained 9.3% of the overall variance in brain temperature, of which 7.6% was redundant with the contribution of the other two factors. Finally, we found that the model is able to highly reliably predict brain temperatures in mice solely based on the sleep-wake distribution and the parameter settings obtained in the other cohort of mice. With our model approach we document and quantify the fundamental dependence of brain temperature on sleep-wake state.

## Results

Wakefulness and REM sleep oppose NREM sleep in terms of brain temperature dynamics. In the cortical activated states wakefulness and REM sleep, brain temperature increases, while during NREM sleep when cortical input is reduced and neuronal activity becomes synchronized, temperature decreases with time (Obal et al., 1985; Franken et al., 1992a). Therefore, based on a previously described model in the rat (Franken et al., 1992b), we used the following exponential equations to iteratively simulate changes of brain temperature in the mouse:

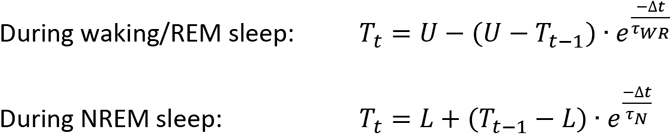

With a time step (Δ*t*) of 0.0011 hours (i.e., the 4-second epochs at which sleep-wake states were scored), the current temperature (*T*_*t*_) is calculated based on the preceding temperature (*T*_*t*−1_) according to the distance from an upper asymptote (U) and time constant τ_*WR*_ when the mouse is awake or in REM sleep at time *t* or, when in NREM sleep, according to the distance from the lower asymptote (L) and time constant τ_*N*_. The four free parameters of this basic model include the two time constants (in hours) and the values of the two asymptotes (in °C) between which brain temperature can vary. Relying on past results and assumptions (Franken et al., 1992b), we initially set the values for both time constants to 0.47 hour, for the lower and upper asymptotes to the minimum and maximum temperatures reached in each animal during the 96 hour recording, respectively, and initial temperature (*T*_0_) was calculated as the average temperature in the first five minutes of recording. The following features were further developed in order to improve the simulation over that earlier study: 1) since representing the two distinctive temperature dynamics with one time constant is likely to be an oversimplification, we allowed different time constants for the increase and decrease in temperature, 2) the asymptote values were free parameters, 3) all free parameters were simultaneously optimized for each mouse individually, 4) optimization took into consideration the entire recording, including sleep deprivation and recovery, instead of only the baseline period, and 5) performance was assessed at a sub-minute scale; i.e., the 4-second resolution at which the sleep-wake states were determined, rather than at an hourly resolution.

With this more elaborate model (i.e. Model0) we could closely simulate the temperature recordings based on the individual sleep-wake state sequence information (Figure 1), with an average correlation coefficient (r) of 0.91 and a median root mean square (RMS) error of 0.36°C. The median time constant of wakefulness was 0.33 hour and usually (i.e., in 9 out of 11 mice) higher than the one of NREM sleep (0.23 hour), indicating that the rate of increase of cortical temperature during wakefulness is slower than its decrease in NREM sleep. The difference between upper and lower asymptotes varied around 3°C, with the upper asymptotes consistently near the temperature values observed during sleep deprivation (Supplementary-Table 1).

**Figure 1:**
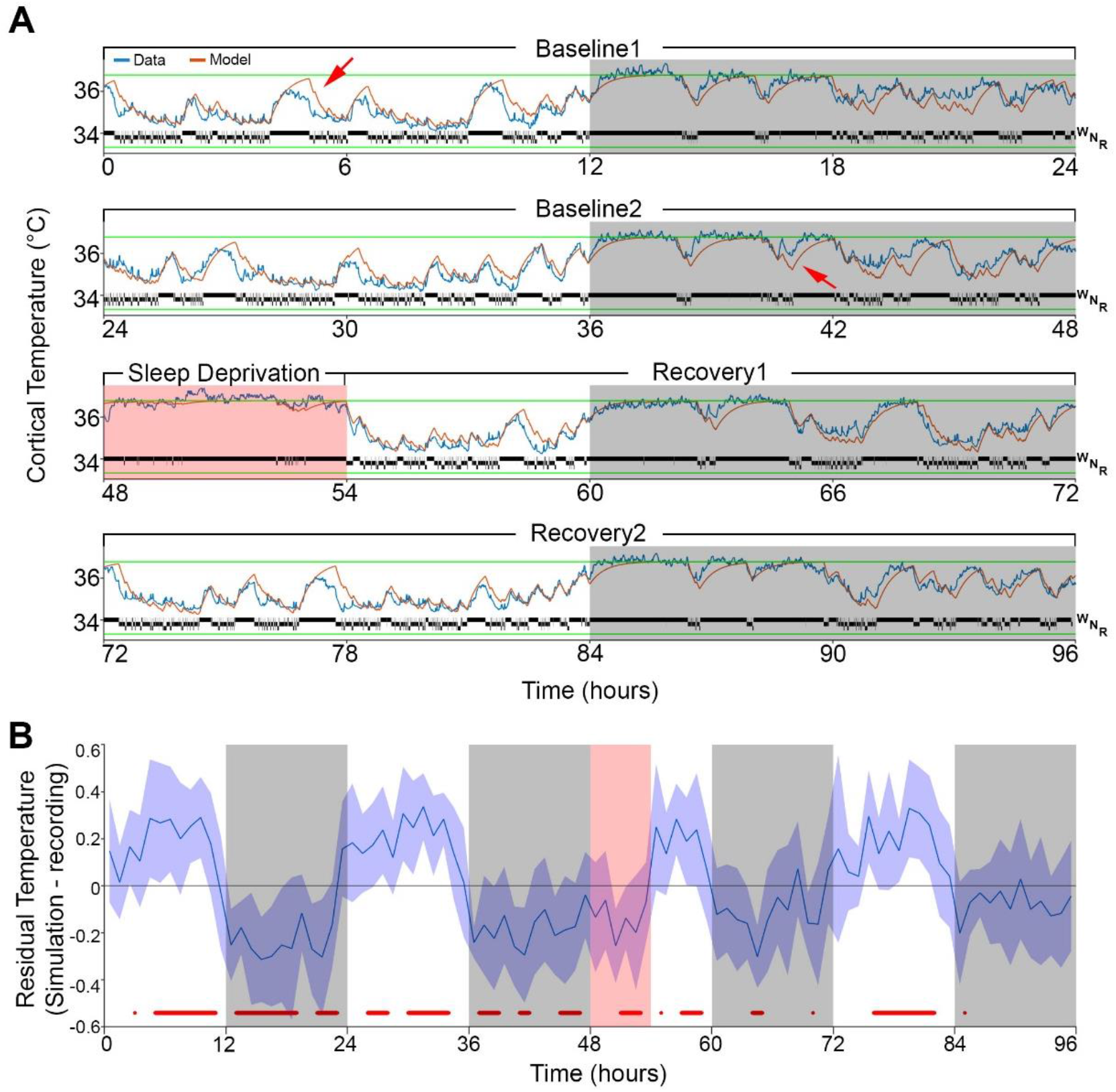
Results of Model0 with constant asymptotes. **A)** Fit example of basic model to raw data. A representative example of 96-hour recording in one mouse of brain temperature (blue) and simulated data (orange). Green lines represent the model lower and upper asymptotes. Above the lower asymptote appears the 4-second hypnogram of wake (W) NREM sleep (N) and REM sleep (R). White/gray backgrounds represent 12-hour light/dark periods respectively, while salmon background starting at time 48 hours indicates the 6 hours of sleep deprivation. Red arrows point to examples of over/under estimation of the model in the light/dark period respectively. **B)** Hourly differences (mean ± STD) between simulation output and data. Median residual RMS across animals and hours is 0.19°C. Red marks below the graph represent significant differences, tested by paired t-test and FDR corrected at p<0.05. Hourly values are plotted at interval midpoint. White/gray/salmon backgrounds as in panel A.

The simulation showed, however, noticeably deviations from data especially when higher temperatures were reached during the light phase and sometimes around lower values during the dark phase (see arrows in Figure 1A). When we examined the time course of average hourly residuals of the model across animals (Figure 1B), we found a systematic fluctuation in the fit with baseline simulated values being too high in the light and too low in the dark phases compared to the recorded temperature values, with a residual RMS of 0.19°C. However, the residuals during sleep deprivation, which occurred during the light period, resembled those of the dark period, arguing against a simple circadian modulation. Indeed, although incorporating a circadian modulation of both asymptotes as previously done in the rat (Franken et al., 1992b) somewhat improved the overall fit (RMS error = 0.32°C; residual RMS = 0.15°C), it didn’t abolish the apparent periodicity in the baseline residuals in mice and resulted in an even poorer fit during sleep deprivation (Supplementary Figure 1). Given the time-of-day independent similarity between the residuals during the sleep deprivation and the dark phase, an alternative factor that could contribute to a temporary upregulation of brain temperature (beyond the sleep-wake state driven changes already captured by the simulation) are periods of sustained wakefulness (Obermeyer et al., 1991). We explored this possibility by, instead of a circadian modulation, changing the asymptotes according to the prevalence of wakefulness (and REM sleep, for consistency) prior to each data point. We refer to this factor as ‘prior wake prevalence’. The window size over which the prior wakefulness prevalence was calculated, as well as the time lag with which it affected the asymptotes, were kept as free parameters. Modulation of the asymptotes according to prior wake prevalence considerably improved the fit (r = 0.95; RMS error = 0.28°C; Figure 2) and removed most of the excessive increases previously noted around high values during light periods. We found that the optimal window size was 4.0 hours, with a shift of 1.5 hours prior to the time point under consideration, and a scaling factor of 1.2°C, which represents the maximum possible modulation of either asymptote (i.e., 100% wakefulness or 100% NREM sleep during a given 4.0h window; Supplementary-Table 2). We optimized all the parameters simultaneously in this new model (i.e. Model1) and found that all of the values obtained with Model0 were changed (all p values ≤ 0.002; F(2,10) ≥ 9.98, 1-way rANOVA); the time constants were almost cut in half to 0.18 and 0.13 hour for wake/REM and NREM sleep, respectively, and the lower asymptote increased leading to a substantial reduction of the inter-asymptote temperature range (from 3.1 to 2.0°C).

**Figure 2:**
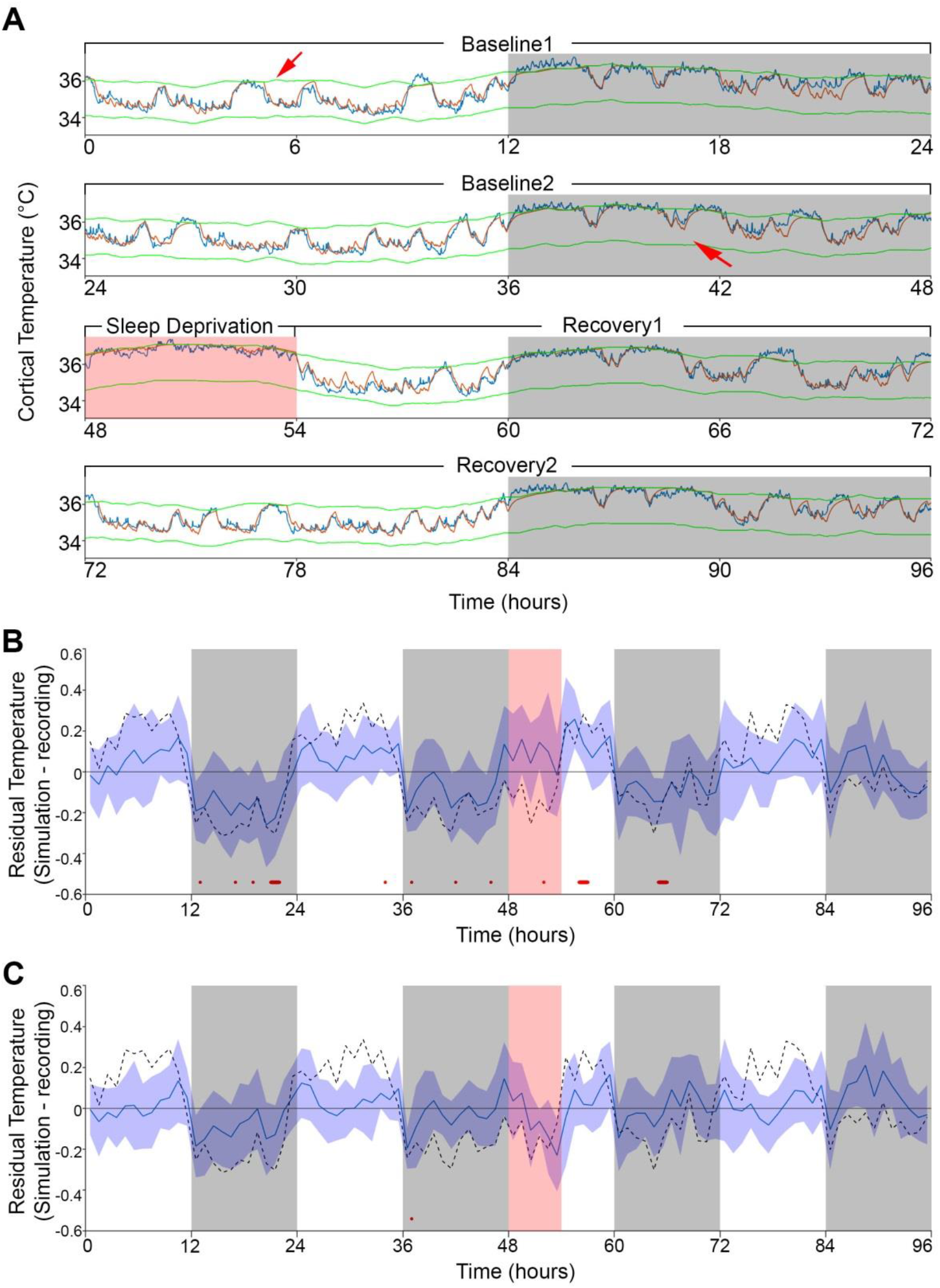
Results of Model1 and −2 with modulation of asymptotes. **A)** Simulation fit after incorporating prior wake prevalence. Note that both asymptotes are modulated in parallel, recorded data (blue line) are of the same animal as in Figure 1, and previous over and under estimations marked by red arrows are diminished. Details as in Figure 1A. **B-C**) Residuals of Model1 and −2, as in figure 1B, after addition of prior wake prevalence (B, Model1) and with an additional circadian rhythm modulation of both asymptotes (C, Model2). Note the reduction in number of red markers indicating significant deviations from zero, with a residual RMS of 0.12°C and of 0.09°C, respectively. Dashed line marks the Model0 mean residuals from Figure 1B.

Interestingly, after incorporating the prior wake prevalence factor, the residuals of the new model now showed a consistent light-dark (circadian) modulation (Figure 2B), with an over-estimation of temperatures in the light periods, including the sleep deprivation (residual RMS = 0.12°C). To account for those errors, we then integrated again a 24-hour sine-wave modulation onto the asymptotes. The combined effect of modulating the asymptotes according to prior wake prevalence and to circadian time (i.e. Model2) almost flattened the residuals (residual RMS = 0.09°C; Figure 2C) and further improved the fit (r = 0.96; RMS error = 0.26°C). Sine-wave amplitude was 0.19°C with a phase of −0.63 hours (placing the trough of the circadian influence at ZT5.37), while window size over which prior wake prevalence was calculated reduced to 3.0 hours and difference between asymptotes reduced as well (to 1.85°C; Table 1). Relative to Model1, the only free parameter that showed a significant change was the scaling factor of the prior wake prevalence window (reduced to 1.01°C, paired t-test, T(10) = 5.4; p = 0.0003). Figure 3 shows the final fit of Model2 to the temperature data for each of the 11 animals. Note mouse #616 had exceptionally long waking periods (see periods of uninterrupted high temperature levels in the dark periods) that likely contributed to its exceptional long prior wake prevalence window (5.5h) and the high ratio between the scale factor (1.05°C) circadian influence (0.11°C), and might explain the aberrant phase of the trough of the circadian factor at (ZT15.6).

**Figure 3:**
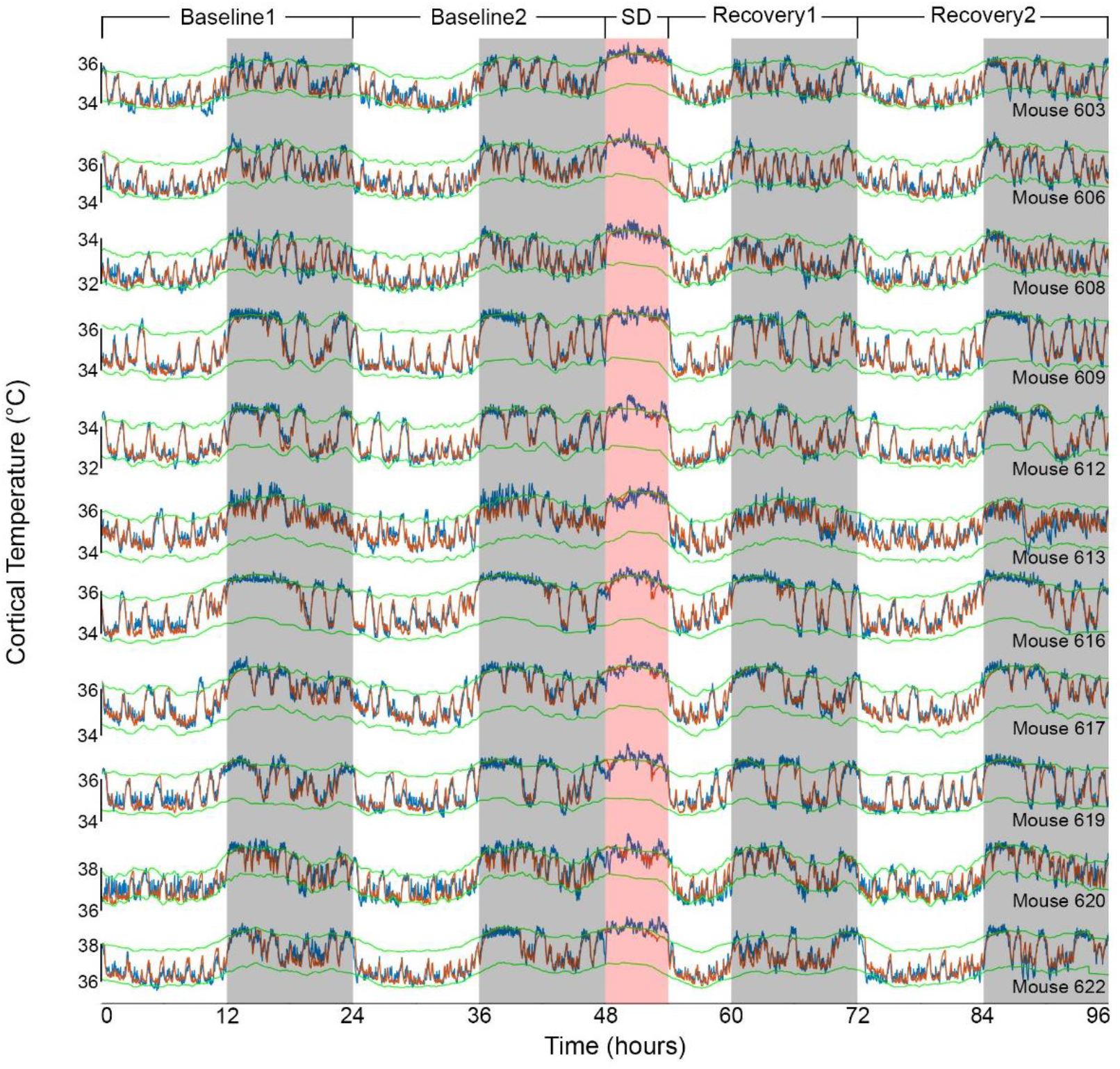
Results of Model2 for all individual mice. The graphs show the fit of the final model for all 11 animals (ordered as in Table 1). Example data in Figure 1A and 2A correspond to mouse number 617 in the 8^th^ row. Color coding as in Figure 1A.

**Table 1:** Model2 parameters for each animal. The table shows the optimized values for each of the parameters, and additional three descriptive variables: the difference between asymptotes (4^th^ column), and root mean squared error and correlation coefficient (two last columns). Columns 7-9^th^ stand for the window parameters of prior wake-prevalence; Window size which is the duration the algorithm took into account, shift is the gap between window ending and the time point been evaluated, and the scale is twice the amplitude of the modulation signal calculated. Columns 11^th^ is the circadian phase, shifting the 24h sinewave (starting at value of zero) relative to ZT0. Last row summarizes median value for each parameter, except the correlation coefficient that is averaged with Fisher transformation. Animal number marked with an asterisk indicate a KO mouse.

| Animal | Asymptotes (°C) |  |  | Time Constants (h) |  | Prior Wake-Prevalence |  |  | Circadian |  | RMS Error (°C) | Correlation |
| --- | --- | --- | --- | --- | --- | --- | --- | --- | --- | --- | --- | --- |
|  | Lower | Upper | Difference | Wake/REM | NREM | Size (h) | Shift (h) | Scale (°C) | Amplitude (°C) | Phase (h) |  |  |
| 603 | 34.26 | 35.82 | 1.56 | 0.21 | 0.06 | 4.75 | -1.9 | 1.02 | 0.13 | 0.86 | 0.28 | 0.94 |
| 606 | 34.74 | 36.59 | 1.85 | 0.23 | 0.1 | 2.75 | -1.3 | 1.01 | 0.25 | -0.71 | 0.26 | 0.95 |
| 608 | 32.28 | 33.77 | 1.49 | 0.22 | 0.08 | 2.75 | -1.4 | 0.92 | 0.25 | -0.41 | 0.23 | 0.95 |
| 609 | 34.04 | 36.28 | 2.24 | 0.22 | 0.11 | 1.75 | -1.1 | 0.68 | 0.19 | -2.66 | 0.25 | 0.97 |
| 612 | 32.6 | 34.45 | 1.85 | 0.23 | 0.11 | 1.5 | -0.9 | 0.73 | 0.18 | -0.63 | 0.24 | 0.97 |
| 613 | 34.15 | 36.07 | 1.92 | 0.11 | 0.17 | 3.25 | -1 | 1.12 | 0.26 | -0.83 | 0.3 | 0.93 |
| 616* | 34.01 | 36.11 | 2.1 | 0.17 | 0.14 | 5.5 | -2.7 | 1.05 | 0.11 | 9.59 | 0.26 | 0.97 |
| 617* | 34.54 | 36.3 | 1.76 | 0.16 | 0.16 | 3.25 | -1 | 1.01 | 0.17 | -1.17 | 0.23 | 0.96 |
| 619* | 34.53 | 36.4 | 1.87 | 0.21 | 0.11 | 3 | -1.6 | 0.62 | 0.06 | -1.07 | 0.25 | 0.96 |
| 620* | 36.89 | 38.21 | 1.32 | 0.12 | 0.05 | 2.75 | -1.4 | 1.16 | 0.22 | 1.29 | 0.29 | 0.95 |
| 622 | 36.14 | 38.11 | 1.97 | 0.2 | 0.09 | 4.75 | -2 | 1.02 | 0.19 | -0.41 | 0.31 | 0.95 |
| Median | 34.26 | 36.28 | 1.85 | 0.21 | 0.11 | 3 | -1.4 | 1.01 | 0.19 | -0.63 | 0.26 | 0.96 |

Since Model2 explained around 91% of the variance in the data, we wondered to what extent each of three factors contributed (Supplementary Figure 2). For this, we disassembled the simulated temperature signal into its 3 constituent factors by removing the circadian and/or prior wake prevalence factors, and subtracting the respective result from the model2 output (see Table 1 and Methods). Consistent with a R^2^ value of 83% reached in Model0, the factor sleep-wake state accounted for the largest portion of the variance; 74%. Some of this explained variance was, however, shared with the other two factors in the model; i.e., prior wake prevalence (27%) and circadian (4%) factors, leaving 42% as its unique contribution. In comparison, the uniquely explained variance of prior wake prevalence (12%) and circadian process (2%; with a 3% shared variance between them) were considerably smaller.

In 5 of these 11 mice, we ran additional experiments with similar design but shorter sleep deprivation (2 and/or 4 hours, starting at ZT0) and only 1 day of recovery. To verify whether the parameters found in the main experiment were not overfitted to the specific experiment, we tested the performance of Model2 in each of the additional recordings, using the individually optimized parameters found previously in the 6h SD experiment (Table 1). Simulation proved to be robust in all the new recordings (all r values ≥ 0.91, and RMS errors ≤ 0.37), convincingly demonstrating the generalization of parameters within animals across different experiments (Supplementary Table 3).

Finally, due to the surprising precision of the simulations, we wondered whether the algorithm can go beyond simulations based on parameters adjusted specifically to an individual mouse. To this end, we used the median parameters values found previously (Table 1) in order to predict the brain temperature of additional mice group (n=5) recorded in the context of another study. The experiment had the same 96-hour design as the current study, and none of those mice data were used to optimize the model parameters. To test the model without providing temperature data, and because the model is iterative, we needed to estimate the initial temperature (*T_0_*) to start the first iteration. We estimated *T_0_* for each animal based on the percentage of wake/REM state in the first 7 minutes of the recording according to the linear regression associating the two variables in the main dataset (see Methods, and Supplementary-Figure 3). Remarkably, all correlation coefficients were between 0.93 and 0.95, although median RMS error was 0.48°C due to some recordings having consistent differences in absolute temperature from physiological values as was also observed in the main data set (see Table 1). When we brought the empirical temperature data to the same average level as the predicted temperature traces (without changing scale), the median RMS error was reduced to 0.34°C and the fine overlap between predicted and observed temperature measures was again revealed (Figure 4, and Supplementary Figure 4).

**Figure 4:**
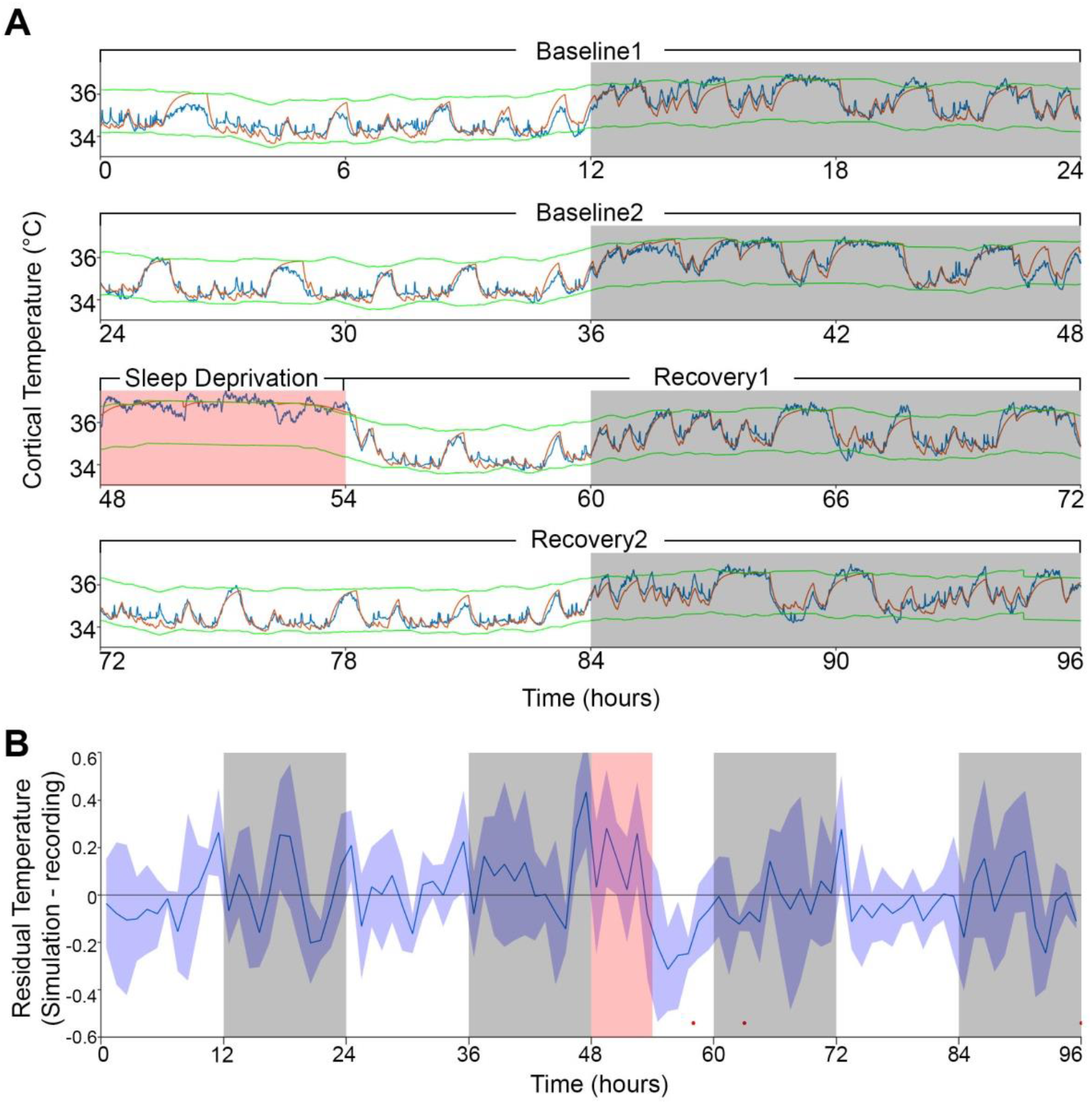
Model fit to a novel dataset. **A)** Representative example of the Model2 fit to a novel raw data not used for optimization, using the median of the optimized parameters from original dataset (Table 1). **B)** Residuals of Model2. Both panels’ details are as in Figure 1A and 1B respectively.

## Discussion

We developed and made available a tool to predict brain-temperature dynamics based on the sleep-wake state sequence. The model showed a remarkable accurate fit with data obtained both under undisturbed baseline conditions and during and following sleep-deprivation challenges of varying lengths. It equally well predicted the global temperature dynamics at a time scale of hours and the changes following sleep-wake transitions in the order of seconds. In addition to two known factors modulating brain temperature; i.e., sleep-wake state and circadian influences, we discovered a not previously recognized contributing factor involving the prior wake prevalence which accounted for the up-regulation of brain temperature observed during the periods of sustained wakefulness.

### Model parameters

In mice kept under our experimental conditions – specifically a 25°C ambient temperature, a 12h:12h light-dark cycle, singly housed, and *ad libitum* food – we observed a dynamic brain-temperature range of a little over 3°C. However, at any given time of the experiment, the range of observed temperatures did not surpass 2°C. This range, delimited by the upper and lower asymptotes in the model, represents a homeostatically defended range within which brain temperature can vary according to the animal’s behavior without eliciting a thermoregulatory response (Parmeggiani et al., 1975; Satinoff, 1983). Both asymptotes, in turn, were modulated by two factors. The first was a time-of-day factor modeled according to a sine-wave, lowering and raising the defended temperature range with an amplitude of 0.19°C, reaching lowest levels close to light-period midpoint; i.e., ZT5.4. Although we referred to this factor as ‘circadian’, given the experimental conditions, this fluctuation could also relate to the imposed light-dark cycle. Nevertheless, using a different analytical approach, our previous work in rats kept under different photoperiods and under constant-dark conditions arrived at very similar amplitudes for the time-of-day modulation of cortical temperature (0.13-0.21°C; Franken et al. 1992b, 1995), supporting the interpretation that the time-of-day factor does not depend on lighting condition and is likely to represent a modulation of circadian origin.

The model identified a second factor modulating the limits of the defended temperature range which we referred to as ‘prior wake prevalence’. This factor quantifies an up-regulation of the level at which temperature is regulated after sustained periods of wakefulness. Up-regulation of brain temperature during sustained wakefulness has been reported on previously but was observed under conditions of sleep deprivations spanning several weeks (Obermeyer et al., 1991). Because in our study the prior wake prevalence effect was transient, concerning a 3h time window only, and was equally observed under baseline and sleep-deprivation conditions, the underlying mechanisms might not be the same. Interestingly, the prior wake prevalence effect was not immediate and affected asymptotes 1.4 hours later. Although we don’t have a good explanation for this finding, delayed effects of sleep-wake state transitions have been reported in humans for body temperature (Bunnell et al., 1988; Youngstedt et al., 1997), suggesting that the delay we observed in the mouse might relate to lagging peripheral effects affecting body temperature and, subsequently, brain temperature. Although we refer to this factor as prior wake prevalence, it is equally plausible that the predominance of NREM sleep in a given interval drives the modulation. The onset of NREM sleep is known to be associated by an active down-regulation of brain temperature involving peripheral vasodilation concomitant with decreased neural activity (Glotzbach and Heller, 1976). Therefore, 3h time windows during which NREM sleep prevails might elicit a delayed net decrease in brain temperature involving peripheral mechanisms. Our current data do not lend themselves for identification of this factor’s physiological substrate and needs further investigation.

Remaining parameters concern the time constants describing the changes in cortical temperature occurring at sleep-wake transitions. To limit the number of free parameters in the model we did not differentiate between the rate of increase in wakefulness and REMS. Although this assumption is likely to be false (see e.g. Hoekstra et al. 2019), since REM sleep is relatively sparsely represented in our data (6% of recording time), we reasoned it would not importantly impact the model’s performance. The simulation identified time constants in the range of minutes for both the increases during wakefulness (and REM sleep) and the decreases in cortical temperature after NREM sleep onset. It also found longer time constants for the increase than the decrease (13 and 7 min during waking and NREM sleep, respectively), pointing to a slower buildup of heat relative to its dissipation during NREM sleep. Such difference is consistent with the concept that the blood perfusing the brain acts as heat sink and increases in temperature underestimate the rate of heat production as cooling occurs simultaneously (Hayward and Baker, 1969).

### Variance explained

Using linear regression analysis between hourly values of wakefulness and brain temperature, sleep-wake state was reported to explain 84% of the variance in brain temperature (Franken et al., 1992b; Hoekstra et al., 2019). Although the overall fraction of the variance explained in the final iteration of our simulation (Model2) was higher (91%), the factor sleep-wake state explained less (74%; see Supplementary Figure 2 for a summary). This discrepancy is due to the additional factors the model identified, primarily prior wake prevalence with which the sleep-wake state factor shared an important part of the variance explained. The linear regression results mentioned above are therefore better comparable to the results of Model0, in which only sleep-wake state sequence was considered. Accordingly, Model0 explained a very similar portion of the variance; i.e. 83%. As suggested previously (Franken et al., 1992b), the circadian factor carried surprisingly little information on brain temperature. Of the 9% of the variance that could be attributed to a circadian modulation, most of it was redundant with that of the other two factors and only <2% was uniquely circadian. In the model, both the circadian and prior wake prevalence factors asserted their influence on brain temperature by modulating the asymptotes. Consistent with the 5-fold larger portion of explained by the latter, also the maximum possible modulation of the asymptote was 5-fold larger (1.01°C versus 0.19°C).

### Comparison to previous models

Compared to our earlier attempt to simulate brain temperature in the rat (Franken et al., 1992b), we made a number of important improvements. In the previous model only the time-constants were optimized while the estimates for the asymptotes were directly taken from the data. Moreover, the same time constants were used to simulate the increases and decreases of temperature at sleep-wake state transitions. Our current analysis shows that increase and decay rates importantly differed with a faster temperature decay in NREM sleep than increase in wakefulness. Additional factors were estimated from the residuals and not formally optimized in one model simultaneously. A first such factor was a time-of-day (‘circadian’) modulation, the amplitude of which was surprisingly similar in the two studies and species (0.15 and 0.19°C, in the rat and mouse, respectively). In the rat study, a down-regulation of the asymptote was needed to accurately fit the initial 12h of recovery because the actual temperature levels were lower than those predicted based on the sleep-wake distribution. Such down-regulation after sleep deprivation was not observed in the current study even without incorporating the prior wake prevalence factor (Supplementary figure 1). This difference could reflect a thermoregulatory response to the longer sleep deprivation in the rat (24h versus 6h in the mouse) or, alternatively, result from the assumption of equal decay and increase rates, because during recovery time spent in NREM sleep is increased which in the current study was found to be associated with a faster decay rate. A final difference between the two studies is the considerable slower time constants in the rat (0.49h) compared to the mouse, a difference that could be attributed to a species difference.

Differences between the simulations were probably also affected by the changes made to address previous criticism. As previously noted by others (e.g. Witting and Mirmiran 1997), sequential evaluation of the contribution of different factors may lead to an overestimation of those assessed first because shared (or redundant) variance will be added to the first but not the second. We addressed this problem by optimizing the parameters for the various factors simultaneously and by explicitly assessing the shared variance among factors. In addition, filtering the data by averaging over hourly intervals removes a large portion of the variability leading to inflated correlations (Heller et al., 2011). We circumvented this drawback by evaluating the performance of the model at the resolution at which the sleep-wake states were assessed (i.e., 4-seconds), which is a more appropriate time scale to study brain temperature changes following neuronal activation (Kiyatkin et al., 2002).

## Conclusions

As the high performance of the simulation could be generalized beyond a specific cohort of mice, a number of applications can be envisioned. For instance, studies assessing the effects of psychoactive drugs on brain temperature (Kiyatkin, 2018), will be able to evaluate and separate direct effects on temperature from indirect effects caused by eventual drug-induced changes in sleep-wake state. Even if there is no effect on sleep-wake state, by recording sleep-wake states those studies, using the model, will be able to generate expected temperature dynamics that then can be contrasted directly to the empirical temperature to assess better the effect of the drug within subjects. Brain temperature affects many properties of neuronal functioning (Kiyatkin, 2010) and therefore may impose critical modulations on cognitive performance (Walter and Carraretto, 2016). Since the dynamics of brain temperature have now been formulated by the model, it may be interesting to examine associations between impaired cognitive functioning and distinctive thermoregulatory states, as happens in sleep inertia just after awakening (Krauchi et al., 2004). However, prior to this, it has to be determined whether brain temperature in humans underlie similar rules. Although such data in humans are sparse, one study reported similar temperature dynamics to those in the mouse (Landolt et al., 1995). The authors, however, reasoned that the underlying driving influence is a circadian rhythm rather than the sleep-wake state, based on locomotor activity and sleep depth arguments. Given that later studies in rodents showed that both activity (Shirey et al., 2015; Hoekstra et al., 2019) and sleep intensity (Franken et al., 1991; Tobler et al., 1994) has only minimal association with brain temperature, it is likely that also in humans brain temperature might be driven by the sleep-wake distribution, in line with the assumptions of our simulation.

## Methods

### Data Acquisition

Detailed description of data acquisition, surgical procedures and experimental design can be found elsewhere (Mang and Franken, 2012; Hoekstra et al., 2019) and will be briefly described below. Data from 11 male C57BL6/J mice (7 wild-types (WT) and 4 lacking the gene encoding cold-inducible RNA-binding protein (*Cirbp* KO mice), 10-15 weeks of age were included. Sleep-wake distribution and brain temperature were unaltered in KO mice (Hoekstra et al., 2019) and the results reported here likewise did not statistically differ between the two lines of mice. All mice were housed individually under a 12:12 hour light-dark cycle with Zeitgeber time (ZT)0 and ZT12, referring to light onset and dark onset, respectively. Ambient temperature was maintained at 25°C and food and water were provided *ad libitum*. Electroencephalogram (EEG), recorded from a frontal-parietal derivation, and electromyogram (EMG), recorded from the neck muscles, were used to ‘score’ the sleep-wake states ‘wakefulness’, ‘NREM sleep’ and ‘REM sleep’, at 4-second resolution. Sleep-wake states marked as having EEG artefacts were included in the temperature analyses. Brain temperature was measured by a thermistor that was placed on top of the right visual cortex corresponding to the midpoint of the front-parietal EEG electrode pair on the left hemisphere, and was sampled at 10 Hz and the median value for every 4-second epoch was calculated. Recordings lasted 96 hours, started at light onset, and included: two days of 24 hours serving as baseline (termed baseline1 and baseline2), 6 hours of sleep deprivation (SD) starting at light onset of day 3, followed by an 18-(recovery1) and a 24-hour (recovery2) recovery period. Sleep deprivation was achieved by gentle handling (Mang and Franken, 2012). In addition to the main experiment above, a subset of 5 animals (1 WT and 4 KO mice) was subjected to a 2- and/or 4-hour SD following the same protocol as for the 6 hour SD experiment but without recovery2. A final, an independent cohort of 5 mice of the same strain, same sex, same age and underwent the same 4-day protocol with 6h SD, was used to test the predictions of the model. All experiments were approved by the Ethical Committee of the State of Vaud Veterinary Office Switzerland under license VD2743 and VD3201.

### Analysis

#### Model0: Optimization details

All code was programmed in Matlab and optimization was done using the *fmincon* function of the Optimization Toolbox, by minimizing the mean squared error between the simulated and recorded temperature signal. All free parameters were always optimized simultaneously, even when additional ones were added at later stages. For each animal we constrained the base values of both asymptotes, to the range of empirical temperature values plus two °C deviation in either direction. The upper and lower asymptotes were always defined as vectors with same size (86’400 4-second epochs) as the 96-hour experiment recordings. The code assumes 12:12 light/dark cycles, start of recording at light onset, sleep scoring includes only wake, NREM sleep and REM sleep states, and 48 hours of baseline.

#### Model1: Modulation of asymptotes according to prevailing state

To modulate the asymptotes in each time point based on the preceding prevalence of sleep-wake state (referred to as ‘prior wake prevalence’), for each time t, we calculated the fraction (0-1) of time spent in wake or REM sleep within a given time window ending at time t-1. We estimated temperature values for time points prior to the start of the recording (i.e. before light onset of Baseline1), by averaging corresponding time points from the two days of baseline. Finally, we subtracted the averaged wake/REM fraction and multiplied by 2 so that values are distributed between ±1, and then multiplied the result by a scale factor to translate the fraction of time spent awake/REM sleep into °C modulation of the asymptote. In addition, to enable the window not to end strictly at time t-1, we also implemented a window shift relative to time t by moving the produced vector back and forth in time. Since this shift was allowed in both directions and we could not predict values after the recovery2 period, we assumed zero values (i.e. no modulation of the asymptotes). The outcome vector was added to both asymptotes.

In contrast to the window scale factor and all other free parameters in this study, which are continuous variables, the window size and window shift are discrete parameters that eventually translated into integer numbers of specific cell indexes in a vector. Due to the requirement of the *fmincon* function for differentiability, to optimize those two parameters we applied a method of brute force: (1) we defined possible values for each of the two variables, (2) for each unique combination of the values we ran the *fmincon* on the rest of the parameters and calculated the error, and (3) kept the parameters (continuous and discrete) that yielded the lowest error output. In this study we chose to test for the window size values between 0 to 10 hours (step size of 0.25 hour) and for the window shift values between −5.0 to +0.5 hours (increments of 0.1 hour). For none of the mice was the best fit obtained with parameter values at the limits of those defined ranges.

#### Model2: Addition of circadian modulation of the asymptotes

To introduce a circadian modulation of the asymptotes we used the following formulas:

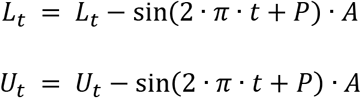

Where *A* and *P* are the free parameters for optimization which stand for the amplitude and phase (in hours) of the sine wave, respectively. *L*_*t*_ and *U*_*t*_ are the lower and upper asymptote values at time t, so both asymptotes were changed in parallel and to the same extent. The minus sign before the sine function is due to the fact that recording started at light onset.

#### Prediction of initial simulated temperature value without actual temperature data

In the original algorithm the initial temperature was determined based on the recorded data, but in order to generalize our algorithm to predict brain temperatures of datasets without thermal recordings we needed a different method to estimate the value of initial temperature. Although temperatures of most animals ranged between 34 and 36°C, some recordings showed non-physiological lower (32-34°C) or higher (36-38°C) ranges (Hoekstra et al., 2019), which probably resulted from technical problems. Nevertheless, the difference between the asymptote values was stable across animals (around 2°C), and simulation performance was high regardless of differences in the average absolute levels in some of the recordings (Table 1). Therefore, to estimate initial temperature using exclusively the state sequence, we produced a predictive formula based on our existing data: First, we normalized the temperature data of each recording in the 96-hour experiment to the range between 34 to 36°C. Then, we determined the correlation between the percentage ‘occurrence’ of wake/REM sleep at a window at start of recording, and the average temperature in the same time window. In this way, we could reliably predict the temperature at the first minutes of recording (r = 0.98, Supplementary Figure 3), which we used as an estimated initial temperature. We chose a 7-minute window size since it yielded the highest correlations across values between 1 to 10 minutes (analysis not shown). Finally, we applied linear regression to obtain the following equation:

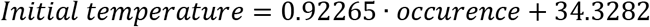

where *occurence* is a value between 0 to 1 with 1 referring to 7 minutes of continuous REM sleep and/or waking, and 0 indicating uninterrupted NREM sleep. Note that this estimate of the initial temperature is valid only for recordings that start at light onset and under a 12:12 hour light-dark cycle.

#### Units and Statistics

Throughout the manuscript time measures are expressed in hours (h), temperature in degrees Celsius (°C), and correlation coefficients (r) are outcomes of Pearson correlations, unless stated otherwise. Optimized parameters are summarized as median values, and correlation coefficients as average after Fisher-transformation (Fisher, 1915; Silver and Dunlap, 1987). Finding significant effects along the time series of 96 hours was done by paired student’s t-tests with false discovery rates (FDR) correction (Benjamini and Hochberg, 1995) for multiple comparisons at a significance level of p<0.05. Individual fits are summarized as the median of root mean square of the difference between simulated data and recorded data (referred to as ‘RMS error’ in text), while effect sizes of the model residuals were assessed by the root mean square of the mean residuals for hourly values (referred to as ‘residual RMS’). Differences in parameters among genotypes and models were assessed by t-tests, or by repeated measures analysis of variance (rANOVA) with Tukey’s range test when more than two values were assessed. The relative contribution of the 3 factors (sleep-wake state, prior wake prevalence, and circadian) to the variance explained by Model2 was calculated as follows; first the model output was disassembled into three traces, corresponding to sleep-wake-state, prior wake prevalence, and circadian. The unique variance explained by each factor was then calculated as the ratio of variance of each factor’s trace and the overall variance in the data. For the shared variance among factors, we subtracted the corresponding unique explained variances from the variance of the traces sum. Results of explained variance are presented with a Venn diagram tool available online (Micallef and Rodgers, 2014).

## Acknowledgements

This study was supported by the Azrieli Foundation (Azrieli Fellowship Award supporting YS), the Swiss National Science Foundation (SNF n°146694 to PF supporting MMBH), and the State of Vaud (supporting MMBH and PF).

## Code availability

The two core Matlab scripts, for brain temperature simulation and parameters optimization, are provided as Supplementary material, together with an example of data and running script.

## Supplementary Tables

**Supplementary Table 1:** Model0 parameters for each animal. Optimized values for each of the four parameters of the Model0 without modulation of asymptotes, and the additional two descriptive variables, as in Table 1.

| Animal | Asymptotes (°C) |  |  | Time Constants (h) |  | Prior Wake-prevalence |  |  | Circadian |  | RMS Error (°C) | Correlation |
| --- | --- | --- | --- | --- | --- | --- | --- | --- | --- | --- | --- | --- |
|  | Lower | Upper | Difference | Wake/REM | NREM | Size (h) | Shift (h) | Scale (°C) | Amplitude(°C) | Phase (h) |  |  |
| 603 | 33.57 | 36.19 | 2.62 | 0.33 | 0.21 | - | - | - | - | - | 0.38 | 0.89 |
| 606 | 33.77 | 37.11 | 3.34 | 0.35 | 0.26 | - | - | - | - | - | 0.39 | 0.89 |
| 608 | 31.49 | 34.2 | 2.71 | 0.33 | 0.23 | - | - | - | - | - | 0.35 | 0.88 |
| 609 | 33.49 | 36.68 | 3.18 | 0.29 | 0.16 | - | - | - | - | - | 0.33 | 0.95 |
| 612 | 31.95 | 34.9 | 2.95 | 0.3 | 0.17 | - | - | - | - | - | 0.33 | 0.94 |
| 613 | 31.81 | 36.78 | 4.97 | 0.36 | 0.81 | - | - | - | - | - | 0.41 | 0.85 |
| 616* | 33.66 | 36.51 | 2.84 | 0.29 | 0.17 | - | - | - | - | - | 0.34 | 0.94 |
| 617* | 33.33 | 36.77 | 3.44 | 0.37 | 0.43 | - | - | - | - | - | 0.36 | 0.9 |
| 619* | 34.32 | 36.7 | 2.38 | 0.3 | 0.12 | - | - | - | - | - | 0.3 | 0.94 |
| 620* | 35.63 | 38.76 | 3.13 | 0.33 | 0.31 | - | - | - | - | - | 0.46 | 0.86 |
| 622 | 34.88 | 38.53 | 3.64 | 0.29 | 0.28 | - | - | - | - | - | 0.41 | 0.91 |
| Median | 33.57 | 36.7 | 3.13 | 0.33 | 0.23 | - | - | - | - | - | 0.36 | 0.91 |

**Supplementary Table 2:** Model1 parameters for each animal. Optimized values of each of the parameters of the model after introducing a modulation of both asymptotes according to the prevalence of Wake and REM sleep in the window preceding the assessment of temperature. Further details as in Table 1.

| Animal | Asymptotes (°C) |  |  | Time Constants (h) |  | Prior Wake-prevalence |  |  | Circadian |  | RMS Error (°C) | Correlation |
| --- | --- | --- | --- | --- | --- | --- | --- | --- | --- | --- | --- | --- |
|  | Lower | Upper | Difference | Wake/REM | NREM | Size (h) | Shift (h) | Scale (°C) | Amplitude (°C) | Phase (h) |  |  |
| 603 | 34.22 | 35.75 | 1.53 | 0.19 | 0.07 | 4.75 | -2 | 1.2 | - | - | 0.29 | 0.94 |
| 606 | 34.55 | 36.48 | 1.93 | 0.19 | 0.13 | 3 | -1.4 | 1.29 | - | - | 0.31 | 0.94 |
| 608 | 32.02 | 33.64 | 1.62 | 0.16 | 0.13 | 3.25 | -1.5 | 1.13 | - | - | 0.28 | 0.92 |
| 609 | 33.78 | 36.37 | 2.57 | 0.22 | 0.13 | 7 | -2.4 | 0.93 | - | - | 0.27 | 0.97 |
| 612 | 32.47 | 34.47 | 1.99 | 0.21 | 0.12 | 2 | -1.1 | 0.86 | - | - | 0.26 | 0.96 |
| 613 | 32.02 | 35.96 | 3.94 | 0.08 | 0.51 | 3.25 | -0.9 | 1.33 | - | - | 0.33 | 0.91 |
| 616* | 33.99 | 36.13 | 2.14 | 0.17 | 0.14 | 5 | -2.9 | 0.92 | - | - | 0.26 | 0.96 |
| 617* | 34.33 | 36.3 | 1.97 | 0.16 | 0.2 | 4 | -1.1 | 1.23 | - | - | 0.25 | 0.95 |
| 619* | 34.43 | 36.5 | 2.06 | 0.24 | 0.11 | 7.25 | -2.7 | 0.78 | - | - | 0.25 | 0.96 |
| 620* | 36.83 | 38.17 | 1.33 | 0.1 | 0.05 | 2.75 | -1.4 | 1.4 | - | - | 0.32 | 0.93 |
| 622 | 36 | 38.05 | 2.06 | 0.18 | 0.12 | 4.75 | -2 | 1.25 | - | - | 0.33 | 0.95 |
| Median | 34.22 | 36.3 | 1.99 | 0.18 | 0.13 | 4 | -1.5 | 1.2 | - | - | 0.28 | 0.95 |

**Supplementary Table 3:**
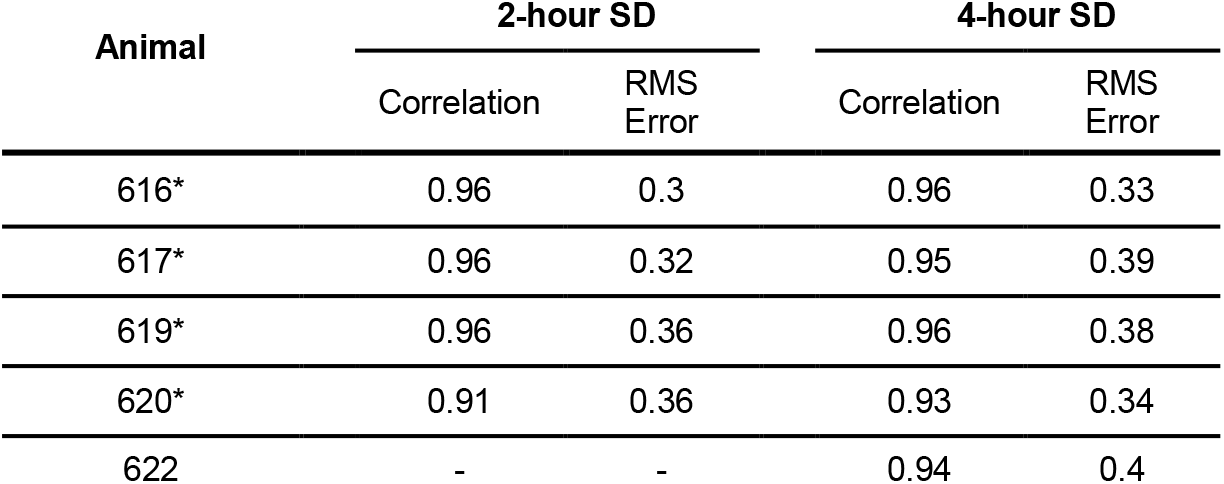
Performance of Model2 for additional SD experiments of the same animals. The table shows the Pearson’s correlation coefficient (r) and root mean squared (RMS) error of five animals from the main experiment, after undergoing additional sleep deprivations (SD) of shorter duration. Due to technical problems the 2-hour SD experiment is missing for mouse number 622. See Table 1 for individual optimized parameters used (asterisks denote KO mice).

## Supplementary Figures

**Supplementary Figure 1:**
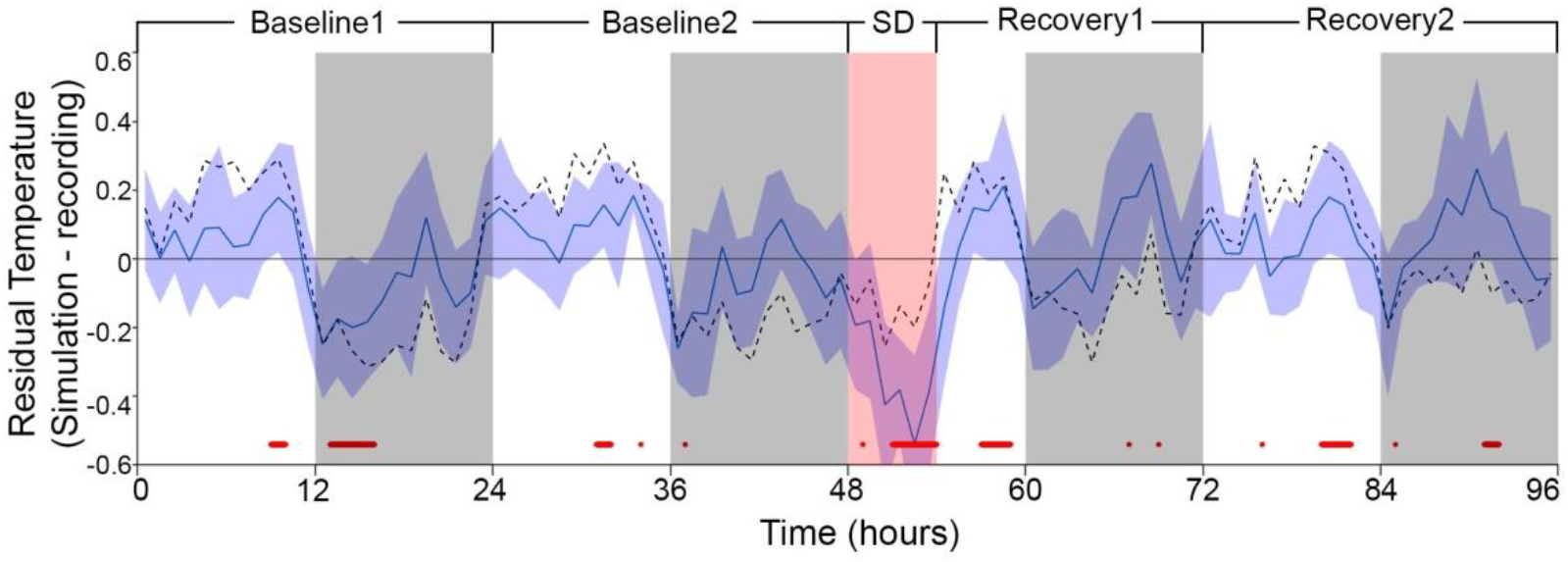
Residuals of the model in which both asymptotes were modulated according to a circadian rhythm. Residuals (as in Figures 1B) of the model after adding a circadian element through modulating both asymptotes according to a sine-wave function with amplitude and phase as free parameters (period was set to 24h). Note the about 0.5°C underestimation during sleep deprivation. Median Residual RMS over all mice and hours amounted to 0.32°C.

**Supplementary Figure 2:**
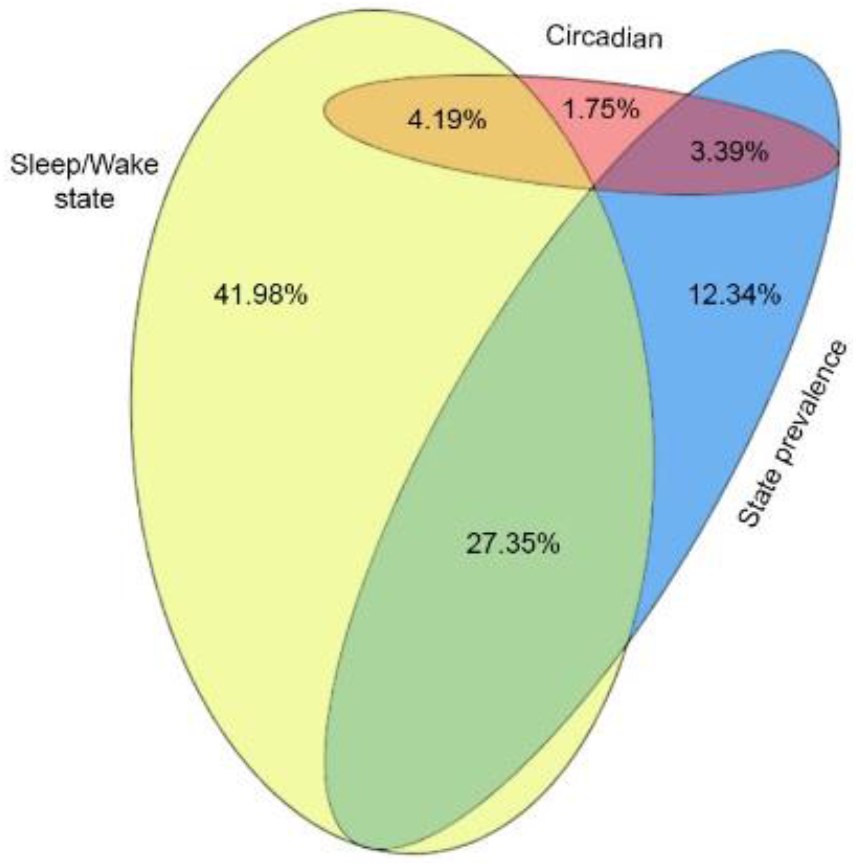
Proportional Venn diagram of the variance explained by each of the three factors in Model2. Yellow (41.98%), blue (12.34%) and red (1.75%) areas correspond to the unique explained variance of sleep-wake state, prior wake prevalence and circadian factors, respectively. The overlapping areas colored in orange (4.19%), green (27.35%), and purple (3.39%) signify the shared explained variance of any two factors. Note the explained variance shared by all three factors was zero.

**Supplementary Figure 3:**
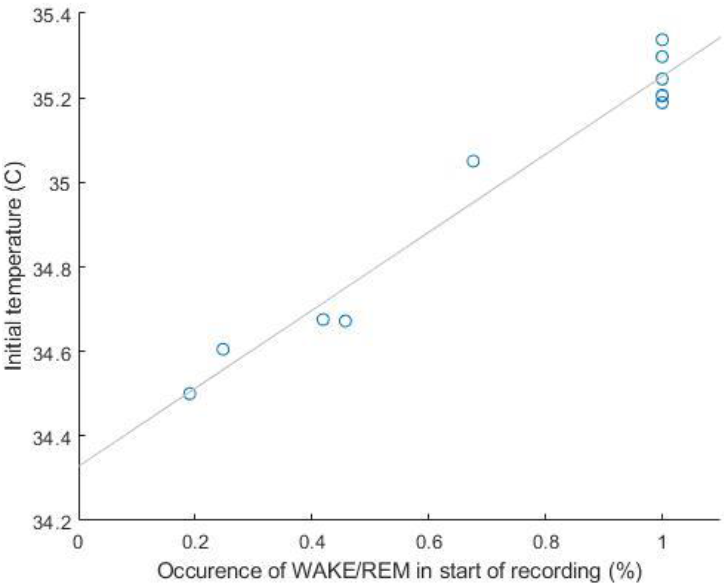
Correlation between initial temperature and Wake and REM sleep prevalence. Average of the normalized temperatures (see Methods) in the first 7 minutes of the recording (i.e., light onset of Baseline 1) is plotted against the fraction occurrence of WAKE/REM sleep in the same time window for the 11 individual recordings from the main experiment. The high correlation allows to predict temperature values of further sessions from sleep-wake state sequence alone.

**Supplementary Figure 4:**
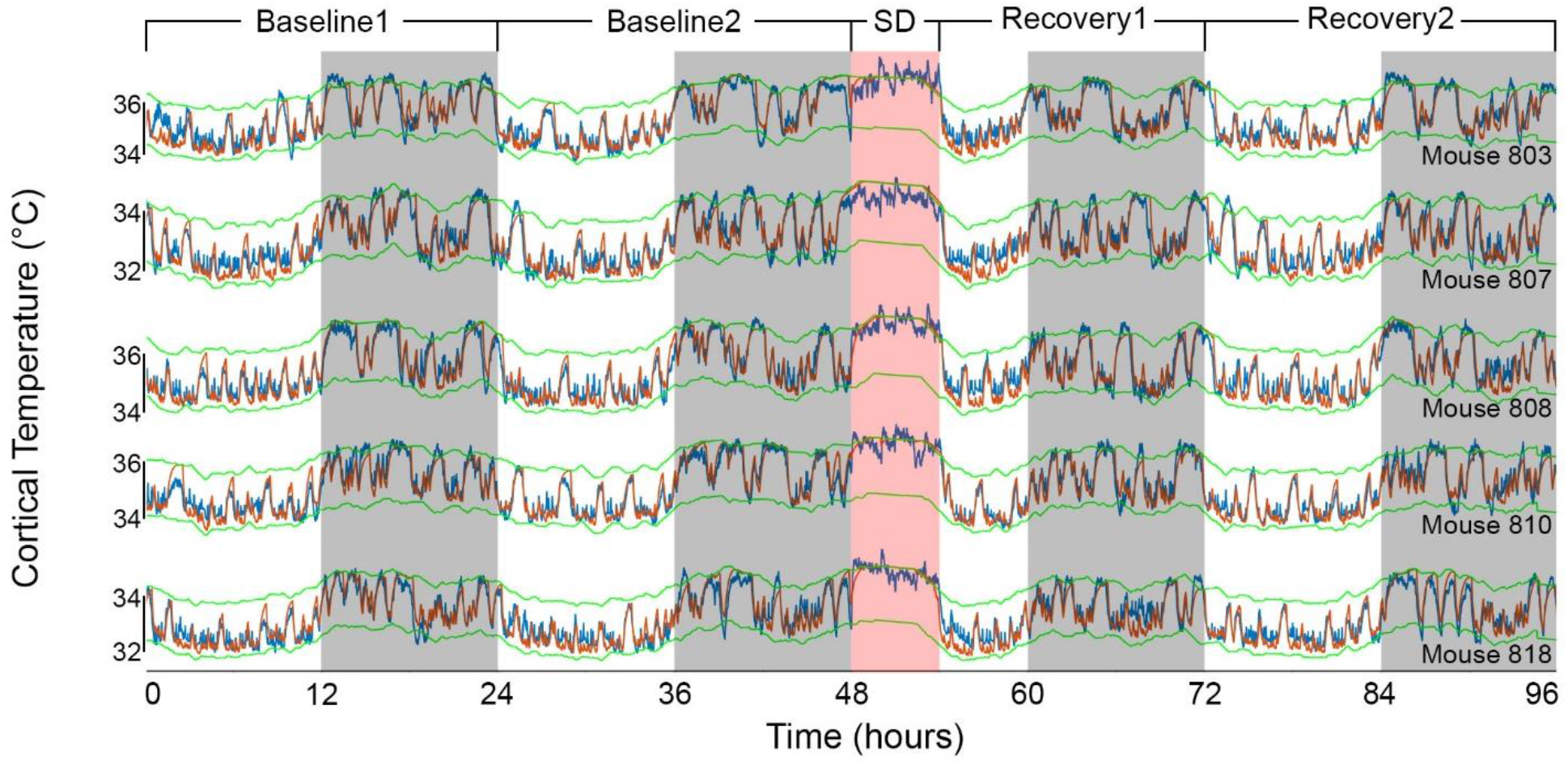
Results of Model2 for independent mice. The graphs show the fit according to Model2, for an additional set of 5 animals, based on the median parameters of our main group of mice (Table 1). Mouse number 810 in the 4^th^ row, appears also in Figure 4A.

